# The McGill-Mouse-Marmoset Platform: A Standardized Approach for High-throughput Imaging of Neuronal Dynamics During Behavior

**DOI:** 10.1101/2020.02.06.937573

**Authors:** Coralie-Anne Mosser, Zeeshan Haqqee, Andres Nieto-Posadas, Keith Murai, Stefano Stifani, Sylvain Williams, Mark P. Brandon

**Author notes:** The authors contributed equally to the work. The authors contributed equally to the work (Sylvain Williams, Mark P. Brandon).

## Abstract

Understanding the rules that govern neuronal dynamics throughout the brain to subserve behavior and cognition remain one of the biggest challenges in neuroscience research. Recent technical advances enable the recording of increasingly larger neuronal populations to produce increasingly more sophisticated datasets. Despite bold and important open-science and data-sharing policies, these datasets tend to include unique data acquisition methods, behavior, and file structures. Discrepancies between experimental protocols present several key challenges including the analysis of the data itself, comparison of data collected between laboratories, and for the comparison of dynamics between brain regions and species. Here, we discuss our recent efforts to create a standardized and high-throughput research platform to address these issues. The McGill-Mouse-Marmoset (M3) platform combines miniscope calcium imaging recording in both mice and marmosets with standardized touchscreen-based behavioral testing. The goal is to curate an open-source and standardized framework for acquiring, analyzing, and accessing high-quality data of the neuronal dynamics that underly cognition throughout the brain in mice, marmosets, and models of disease. We end with a discussion of future developments and a call for users to adopt this standardized approach.

## Goals of the M3 platform

The scope and complexity of data collection in the field of neuroscience, from multiple brain regions at multiple levels of organization, is rapidly growing^1,2^. Technological advances in electrophysiology and neurophotonics have facilitated the mapping of increasingly larger neuronal populations with improved resolution of spiking activity in freely behaving animals^3^. This progress has resulted in rapid dissemination of exciting new insights in network dynamics throughout the brain^4^. As we move away from the analysis of individual neurons to focus on the activity of large networks, we begin to reveal the dynamics and underlying manifolds that govern how activity propagates through neuronal populations to subserve behavior and cognition. Genetically engineered mice have been instrumental in our ability to advance this process with the implementation of advanced genetically encoded calcium indicators (GECIs) to monitor the activity of neuronal populations^5^, along with opto- and chemogenetic approaches to circuit activation and suppression^6,7^.

Despite this substantial recent progress, our understanding of the principles and mechanisms underlying complex brain function and cognition remains incomplete because of three key limitations in current approaches. First, despite rapid progress in recording technologies, the ability to record from all neurons in the brain simultaneously is not a feasible possibility anytime soon. Second, behavioral paradigms vary widely across laboratories, which prevents researchers from efficiently comparing their datasets, leading to what is often referred to as a ‘reproducibility crisis’. Third, to translate our findings beyond the rodent brain, we need to establish these high-yield recording approaches during identical, or at least comparable, behavioral tasks in non-human primates. Together, these limitations hinder our progress in understanding how brain-wide networks and subnetworks interact to support behavior and cognition.

To tackle these challenges, we have established the McGill-Mouse-Marmoset (M3) platform, with the key mandates to (1) provide a blueprint and service to researchers who wish to collect neuronal recordings during behavior, (2) to standardize behavioral tasks during recordings between labs, and (3) to combine recordings across brain regions and species into an open source database to share with the greater neuroscience community.

We offer to substantially accelerate large-scale data acquisition from animal models at behavioral, circuit, and cellular levels by combining cutting-edge calcium imaging techniques with high-throughput standardized behavioral tasks. Calcium imaging involves the use of miniature head-mounted microscopes to monitor the activity of large populations of neurons, as reported by a fluorescent calcium indicator. Recent advances in microscope miniaturization, fluorescent calcium indicators and data analysis pipelines have made possible simultaneous high-fidelity recordings of large populations of neurons (in the hundreds) in freely behaving mice^8^. Such data afford the power to precisely estimate activity at the population level, which is not matched by standard electrophysiological techniques. Moreover, calcium imaging also offers the ability to record chronically on the order of months while tracking individual cells. For standardized behavioral tasks, we use the Bussey-Saksida touchscreen chambers^9^, which are programmable apparatus capable of running more than 14 automated behavioral tests targeting the function of various brain regions in health and disease. The M3 platform has set up 12 of these touchscreens boxes, each equipped for calcium imaging in mice from a miniature head-mounted microscope.

While mice are by far the most widely used models in neuroscience and provide an important fundamental framework for understanding brain function and disease, the translation of basic neuroscience discoveries into disease treatments administered in the clinic has been often hampered by a “model mismatch”, that common rodent models do not always compare effectively to human conditions due to neurophysiological, cognitive, and social differences. We are therefore establishing this same high-throughput approach for recordings in marmosets to facilitate translational research in the near future. The common marmoset *Callithrix jacchus* is one such non-human primate model that overcomes several limitations of current models because of their complex brain anatomy, neural circuitry, behavior, and recent successes in the generation of transgenic marmoset models of human disease. A major advantage of using standardized behavioral tests that are fully automated is that the data will be comparable both between mouse and marmoset experiments, as well as across sites. Moreover, we can adapt cognitive paradigms for mice and marmosets directly from touchscreen-based computerized tests for humans, a crucial factor for the translation of cognitive outcomes across species in preclinical and clinical trials.

Here, we describe the current setup of our apparatus and analysis tools used to collect behavioral and calcium imaging data from mice, with a brief review of miniscope and touchscreen technologies and the advantages that will come with combining these two techniques in neuroscience research.

### Miniature fluorescent microscope imaging to probe neuronal activity

Understanding the neural mechanisms underlying cognitive function requires tools to characterize neural activity from various brain regions *in vivo*. Electrophysiological techniques can probe the activity of multiple neurons with single-cell resolution and high temporal precision^10^, but these approaches are often limited in the maximum number of neurons they can record, as well as their ability to reliably track the same neurons over longer timescales. Thus, such techniques are limited in their ability to resolve population dynamics, which are evident only when large neuronal assemblies are unfolded across days, weeks, or months.

Over the last two decades, many of these limitations have been overcome through the development of optical imaging tools and GECIs^11^, enabling neuroscientists to monitor *in vivo* neuronal activity via microscopy^5^. Imaging techniques for measuring brain activity at both cortical and subcortical levels and at either cellular or subcellular resolutions, such as two-photon^12^ or confocal microscopies^13^, require the animal to be head-fixed, which limits the variety of behaviors that can be tested. With the emergence of complementary metal-oxide-semiconductor (CMOS) sensors, the Schnitzer group developed miniaturized microscopes (miniscope) with epifluorescent light sources^8^. These miniscopes are light, small, and easy to mount on the head of a mouse for direct recording of cell-type-specific activity patterns in freely behaving animals. The GRadient INdex (GRIN) lens, which faithfully relays images from subcortical tissues in the brain to the CMOS sensor, further improved miniscope imaging systems to record from deep brain regions^14^. Such systems allow reliable and consistent recording of calcium activity from hundreds of neurons for months, facilitating the decoding of neuronal representations underlying specific behaviors.

With these advantages in mind, the M3 platform chose to use an open source miniscope known as the UCLA miniscope (www.miniscope.org). This site provides an outstanding open-source platform developed by the laboratories of Peyman Golshani, Alcino Silva, Baljit Khakh, Daniel Aharoni, Tristan Shuman, and Denise Cai for sharing access to all the associated software and hardware files necessary to assemble and implement recordings with this system. The miniscope itself is a flexible, inexpensive and state-of the-art light-weight single-photon miniature microscope designed to detect GECI activity in cells, enabling the imaging of almost any brain region while animals behave freely in tasks such as spatial navigation, complex motor movement, social behaviors, as well as abnormal behaviors in disease models. For our purposes, single-photon miniscopes are currently advantageous over their two-photon homolog due to simpler maintenance and assembly, smaller size and weight, and better affordability^15^.

### The M3’s modified miniscope system

Most procedures used in the M3 platform to assemble a miniscope, such as the insertion of the optics (emission and excitation filters, dichroic mirror, achromatic and half-ball lenses) into the main body, as well as the soldering of the LED connection to the CMOS, are performed following the protocol proposed on the www.miniscope.org website. Nevertheless, we made some modifications to the original UCLA first generation miniscope design to produce a robust, easy to maintain, and long-lasting tool. The miniscope structure is composed of two body parts sliding onto one another to adjust the focus of the image, and is traditionally made of Delrin® acetal homopolymer, known for its malleability and low resistance to repeated contact. Since the behavioral tasks used in the M3 platform require training animals daily over several months, screwing the miniscope to the baseplate risks damaging it. We therefore changed the body part materials to thermoplastic polyetherimide Ultem®, which benefit from excellent strength and stiffness and outstanding stability of physical and mechanical properties at elevated temperatures. Moreover, this resin retains strength and resists stress cracking when exposed to alcohols or acids, which facilitate its cleaning. We also reinforced the wires connecting the CMOS to the LED by wrapping them with heat-shrink tubing, as well as the coaxial cable connecting the miniscope to the data acquisition controller using two springs covered with heat-shrink tubing at each extremity. These modifications considerably reduce the risk of breaking the soldered connections. We also added a custom-made static shield and a heat sink to protect the CMOS and prevent avoid over-heating, as the miniscopes are designed to function continuously for several hours.

The main body of the miniscope houses the optical parts and CMOS imaging sensor, which sends the digital imaging data to a custom data acquisition (DAQ) electronics and USB host controller over a flexible coaxial cable. We modified the DAQ by soldering onto it a transistor-transistor logic (TTL) connection, which allows any external software to trigger the miniscope on or off and synchronize its recording with the recording of the behavioral task, such as inputs from the touchscreen. The data acquisition frame rate of our miniscope is 30 Hz, with 722 × 480 pixels resolution.

For animals implanted with a 1.8mm diameter GRIN lens, we use a regular miniscope setup to record from the brain. For animals implanted with smaller lenses (either 1.0mm or 0.5mm diameters), we insert a 1.8 mm-diameter GRIN lens into the sliding body part to relay the signal from small diameter implanted lenses. Our miniscope dimensions are 2 mm (L) × 1.6 mm (W) × 2.1 mm (H). Its weight is approximately 2.8g (2.9g with the GRIN relay lens) with a 1.2 mm x 0.76 mm maximum field of view (1.0 mm × 0.65 mm with the relay lens).

### Surgical procedure for GRIN lens implantation

Each mouse typically requires three surgeries to prepare for imaging as described in figure 1: (1) viral injection to induce expression of the activity-dependent fluorescent calcium indicator in to-be-recorded areas, (2) GRIN lens implantation above the target brain region, and (3) affixing the miniscope holder. Therefore, it takes approximately 5 to 6 weeks – depending on the viral recombination and expression time course – from the first surgery until the start of data collection. The platform uses adeno-associated viral resources for *in vivo* gene delivery and will soon include several transgenic mouse lines expressing fluorescent reporters in diverse neuronal subpopulations.

**Figure 1.**
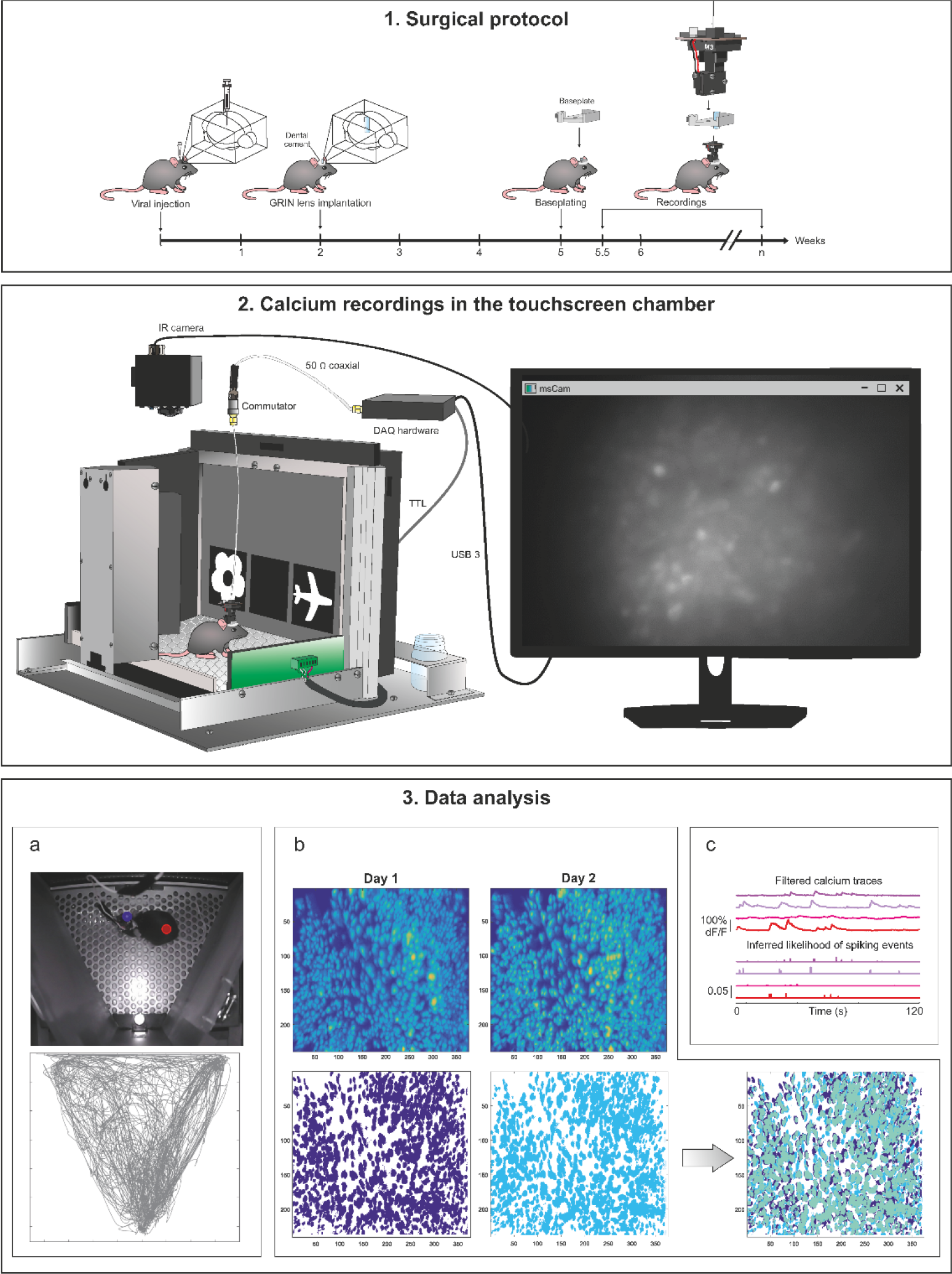
M3 platform workflow. (1) Surgical protocol timeline from viral injection to miniscope recording, with an illustration of the miniscope main body and baseplate. (2) A mouse in a touchscreen chamber completing the PAL task. The miniscope is connected to the computer via a coaxial cable relayed through a multi-channel commutator and a DAQ. The mouse behavior is recorded via an IR camera placed above the chamber. The computer screen depicts real-time video footage of calcium fluorescence recorded during behavior. (3) (a) (Top) A mouse being tracked in a touchscreen chamber using DeepLabCut. Blue dot represents tracking of the mouse body, while red dot represents tracking of the miniscope LED. (Bottom) Path of movement generated from one complete behavioral session. (b) (Top) Spatial footprints of cells extracted from *in vivo* calcium imaging recordings in CA1 for two example sessions (day 1, 1012 neurons; day 2, 1106 neurons). (Bottom) Resulting footprint from the merging of both sessions to track the same neurons across different days. (c) Example of extracted filtered calcium traces (top) and the resulting inferred likelihood of spiking events (bottom).

The GRIN lens implantation is performed two weeks following the viral injection. Our miniscopes can be used for recordings using various GRIN lens size, from 0.5 to 1.8 mm diameter. Inserting large dimeter lenses (e.g. 1.0-to 1.8-mm diameter) requires brain tissue resection to make room for the lens, whereas a 0.5 mm diameter GRIN lens can be directly implanted into the brain after making a leading track with a blunt-end 25G needle. Thinner lenses, while offering a smaller field of view, greatly minimize tissue damage and are thus better suited for imaging deep brain structures. Due to tissue damage along the lens track, a long recovery time (for 3 to 4 weeks) is required before imaging can commence^16^. The final step of fixing the baseplate onto the implanted GRIN lens is not a surgical procedure, but anesthesia is still required to fix the animal’s head in place while cementing the baseplate onto the skull. The baseplate is fixed with the miniscope attached to search for an optimal field of view of and focus. Delicacy in this final step is crucial for the image quality of the cells to be recorded.

### Automated touchscreens operant conditioning chambers

A primary aim of the platform is to reliably translate research between mouse and marmoset studies. Standardized behavioral tasks designed to train and test either species with similar efficiency and that minimize extraneous variables between experimenters will be necessary for a valid comparison of the neural correlates of behavior between these two animal models. The touchscreen operant conditioning chambers developed by Tim Bussey and Lisa Saksida offer numerous compelling advantages to miniscope calcium imaging research across animal models. In addition to greatly reducing experimenter involvement via task automation and minimizing animal stress by training through positive appetitive reinforcement, the Bussey-Saksida touchscreens are equipped with a large variety of behavioral assays designed to target specific cognitive capacities. For instance, in the trial-unique, delayed nonmatching-to-location (TUNL) task, the animals are taught through various stages of training over the course of weeks to remember and compare the locations of pairs of identical visual cues separated by a delay ^17^. By manipulating the proximity of cue locations and the duration of the delay, the task can be made to be increasingly difficult in ways that are thought to assay different cognitive functions (pattern separation/pattern completion and short-term memory, respectively). This task has been shown to be hippocampal-dependent in mice^18^. Another example is the object-location paired-associates learning (PAL) task, which presents pairs of unique stimuli in two of three possible locations on the screen, with one object in its correct location and the distractor in an incorrect location^17^. This task tests location-dependent associative memory ability and is sensitive to hippocampal lesions^19^ and cholinergic muscarinic receptor manipulations in mice^20^. Other tests target the study of attention, visual discrimination, impulsivity, visuomotor conditional learning, Pavlovian conditioning, and location discrimination^21,22,23^. All tasks are also fully reprogrammable on the accompanying ABET II Touch software from Lafayette Instrument®, allowing researchers to manipulate study parameters to cater to any additional variables of interests.

When combined with miniscope calcium imaging and video recordings of behavior, freely behaving animals can be tracked simultaneously in task performance, movement and orientation within the chamber, and neuronal population activity in targeted brain regions can be recorded at the single cell resolution. Recording from different brain regions on a single task enables the creation of a database that maps differences in population activity across the brain for any given cognitive assay, which can then be reliably be compared to the activity mapped in a different animal model. This translation provides a powerful glimpse into whole-brain activity dynamics shared across mice and marmosets and is efficient for validating any findings from disease models in mice to analogous models in marmosets, thereby assisting in our understanding of disease models in humans.

One drawback to using automated touchscreen operant chambers is in slower task acquisition compared to simpler tasks with an optimized apparatus and greater experimenter involvement^21^. This drawback is countered by the ability to run multiple touchscreen chambers simultaneously, allowing researchers to complete daily sessions for a cohort more quickly, as well as conduct multiple different experiments at the same time. Our 12-box setup enables high throughput acquisition of data to accelerate research projects. We also include numerous modifications to the traditional chamber setup to incorporate neurophysiology and video tracking recordings in our analyses.

### Touchscreen apparatus and modifications

Figure 1 depicts a schematic of our setup for a single box, detailing all devices involved in data acquisition during testing and their connections between each other. A high-resolution infrared camera with UV emitters is attached directly above the touchscreen apparatus with its position adjusted using shims to provide an overhead field of view of the platform on which the animal explores during behavioral testing, as well as the stimuli displayed on the touchscreen. The light-sensitive sensor on the camera is blocked using tape to keep it from turning off and on repeatedly during behavioral recording, as a house light often lights up inside the box during certain tasks to indicate an animal’s incorrect attempts.

Each touchscreen task requires the use of a specific mask secured directly in front of the touchscreen, with cutouts corresponding to locations where stimuli are expected to appear. The masks enhance the contrast and visibility of test stimuli by blocking backlight from remaining parts of the touchscreen while also creating distinct tangible spatial locations for the animal to respond to. However, head-mounted implants and miniscopes tend to be taller than the size of the cutouts for some masks, such as those in the TUNL task, making it difficult for the animal to poke its nose through the cutout to touch the screen. As such, we’ve modified some of our masks by making taller cutouts to prevent head-mounted devices from impeding performance.

In the traditional touchscreen setup for mice, the reward port is located on the far side opposite the center of the touchscreen and dispenses strawberry milk reward onto a tray, with an opening for the mouse to poke its nose inside to consume the reward. Large head-mounted implants, such as miniscopes, once again make it difficult for the animal to reach these reward ports. As such, we fixed our apparatus with custom-made reward ports from Lafayette Instrument® that were modified to bring the reward tray outwards with an open top to remove the overhead roof that would otherwise impede animals with head implants from collecting their reward. Bringing the reward tray forward toward the touchscreen also required adjusting the position of the IR beams located on the side of the apparatus, which break when the animal crosses either toward the reward port or toward the touchscreen.

To prevent the animal from jumping out and escaping the chamber during testing, clear plastic lids are fixed atop the touchscreen chamber and secured in place using clamps attached to the side walls of the chamber. For miniscope recordings, our lids have a cutout on the top to allow the coaxial cable attached to the miniscope to connect to the commutator directly above the chamber. For all our recordings, we use multi-channel commutators from Dragonfly Research and Development Inc. that allow us to incorporate electrophysiology with miniscope recordings simultaneously. Our commutators will also soon be compatible for use with fiber optic rotary joints in series, allowing us to include optic fiber recordings alongside miniscope recordings in future experiments.

The coax cable from the miniscope is soldered to the commutator and secured using Kwik-Sil™ and heat shrink wrap. The other end of the commutator is also soldered to a coax cable, which is connected to the DAQ placed on a shelf at the top of the box. Both the DAQ and the infrared camera are connected via USB to a small computer (Intel® NUC Kit NUC8i7BEH) fitted with 16GB of RAM and 1TB of 3D NAND solid-state storage. Data is recorded onto this computer using the miniscope recording software provided by www.miniscope.org. The software enables simultaneous recording of two video streams (miniscope and behavioral video) with a multitude of adjustable video settings for the behavioral recordings (exposure, gain, sharpness, brightness, contrast, etc.) and adjustable exposure, gain, and LED intensity for the miniscope recordings. All recorded data is backed up to cloud servers daily.

Each DAQ in a set of four boxes also connects to the ABET TTL Adapter, provided by Lafayette Instrument®, located outside the touchscreen box. The TTL adapter increases the number of outputs from the main computer to the touchscreens, which is necessary to accommodate all four touchscreen boxes in a rig and provides a direct connection between the individual DAQs and the main computer that controls the four touchscreens, where ABET II Touch software is used to set up and initiate behavioral tasks. In order to utilize the TTL signal to initialize miniscope and behavioral recordings with touchscreen inputs simultaneously, we first add the TTL as a new output device in the environment designer in ABET II Touch. We then use the schedule designer to edit any behavioral schedule of interest to add a condition that turns on the TTL at a specified time point; usually either at the very beginning of the trial or when then animal first initiates the task. The TTL signal can also be set to turn off the miniscope and infrared cameras at a specified time point, after a certain number of trials, or during a specific behavioral event.

A typical session begins with priming the reward tray with strawberry milk reward, delivered via tubing connected at the back of the reward port that mechanically pumps the milk into the tray. After attaching the miniscope to the commutator in one of the boxes, all devices are turned on and checked for proper functioning. An animal is first weighed and then handled carefully to attach the miniscope to the baseplate cemented on its head, secured with a small set screw. After the animal is placed in the chamber, the door compartment is then elevated to close off the entrance and a lid is placed on top, secured and locked in place via clamps on the walls of the chamber. The outer door of the sound-attenuated box is closed to minimize noise disruptions from outside and from the surrounding boxes. The LED intensity and video parameters are set on the miniscope software on the Intel® computer, with the trigger set to turn off the miniscope LED until initiated by the TTL signal. On the ABET II Touch computer, the desired schedule with number of trials and time limit is set and the task is started. Once the animal initiates the task (or whenever the TTL signal was set to activate), the miniscope LED is turned on and begins recording alongside video data. The recordings are terminated automatically when the task ends, or after a set amount of time; typically 20 minutes for miniscope recordings, to minimize photobleaching of the calcium indicator.

### General analysis procedures

The ultimate goal of the platform is to promote dissemination of knowledge and data sharing. The complex data sets to be acquired from mice and marmoset models will require a comprehensive database for data storage and accessibility to users. Currently, the M3 platform is developing a standardized data structure along with pipelines for pre-processing neuronal and behavioral data in order to accelerate verification, reproducibility, and collaboration. General analysis procedures are detailed below and outlined in figure 1.

Following each recording, imaging data is motion corrected^24^, and individual cell data are extracted via constrained nonnegative matrix factorization^25^, generating a graphical representation of overall fluorescence during a recording session. The likelihood of spiking events for each cell is inferred from deconvolution of the calcium trace via a second-order autoregressive model using published preprocessing pipelines^26^. The identity of individual cells is tracked across days to permit cross-session cell-wise and population-wise comparisons^27^. Further analyses are implemented via custom-written code. Mouse position data is inferred from the miniscope LED and the mass center of the mouse body offline using a trained DeepLabCut network^28^. Behavioral data is extracted from the ABET II Touch software in the form of millisecond resolution timestamps of all behavioral and task events that occurred during the selected session, including task initiation, stimulus presentation, stimulus selection, inter-trial-intervals, etc. Simple analyses can be performed correlating behavioral events of interest with patterned spiking of cells, but the primary analysis of interest is the use of generalized linear models (GLMs) to characterize and dissociate contributions of visual cues, spatial location, rewards, and stimulus delays to neuronal and population level representations within and across sessions. Such models are uniquely suited for analyzing data when multiple variables are contributing to the final signal^29^, and these models have previously been successful used in characterize=ing similarly large datasets in other domains of neuroscience, particularly in the analysis of functional magnetic resonance imaging.

### Application and future directions

A popular use for automated touchscreen operant behavioral testing is in mouse models of disease. Many labs have already tested various disease models on a number of touchscreen tasks. These include testing the TgCRND8 mouse model of Alzheimer’s disease on attention, memory, and impulsivity^30^; cognitive impairment in a mouse model of schizophrenia^31^, and reversal learning in a mouse model of fragile X syndrome^32^. Questions that still linger from these studies center around the underlying neurophysiology of the mice during these tasks as well as what this neurophysiology can indicate about the cause of the deficits observed in these animals and, by extension, in the diseases they model. We believe miniscope calcium imaging provides the next natural step in deciphering the population dynamics of the brain regions implicated these diseases to reveal the abnormal patterns of activity that drive their deficits.

Having established a functional protocol for modelling neural correlates of behavior in mice across multiple brain regions, the platform is gradually progressing toward applying the same combination of methods to studying marmosets. Our capacity to achieve this is facilitated through access to the Comparative Medicine and Animal Resources Center (CMARC) at the McIntyre Medical Sciences Building in Montreal, Quebec, as well as the marmoset transgenic core at the Montreal General Hospital. Both institutions will house large experimental breeding colonies of marmosets, with the latter also producing transgenic animal lines. Testing facilities will be situated at the Montreal Neurological Institute (MNI) to make use of its concentrated expertise in non-human primate research, with the plan to incorporate touchscreens as part of home-cage cognitive testing in the animals as they age and develop.

As miniscope technologies continue develop, the platform is expected to evolve with it. Currently, the one-photon, wide-field miniscope is ideal for high throughput imaging in freely behaving animals as it is the most cost-efficient, lightweight, and robust version of the miniscopes developed to date, with a large field of view to maximize cell counts^15^. However, recent initiatives in open source development of miniscopes pave the way for a multitude of independently designed and modified miniscopes in the near future, with a variety of unique uses^33^. These include integration with optogenetic manipulations^34^ and wire-free miniscopes for efficient monitoring of social behaviour^35^. The miniaturization of two-photon microscopy introduces optical sectioning for three-dimensional imaging and improved spatial resolution of cells, which permits tracking of cellular processes at the scale of individual dendritic spines during behavior, while altogether minimizing widespread photobleaching from single-photon imaging^36^. Due to the ‘plug-and-play’ nature of our setup with the automated touchscreen operant chambers, the obstacles involved in updating miniscopes models to include additional data outputs or improved video recording will be limited only by the hardware capacity of our recording devices and computers, as well as modifications to surgical procedures.

Accompanying future miniscope developments is the rapid evolution of neurophotonics. As mentioned earlier, miniscope calcium imaging in mice typically begins long after viral injection to the brain region of interest, as certain promoters take several weeks to become fully expressed. With the use of transgenic lines of GCaMP6f mice, expression of fluorophore is present across the entire brain of the mice from birth, eliminating the need for viral injections and reducing the wait time post-operation to behavior while also minimizing variations in expression strength between animals due to variations in injection efficiency.

Genetically encoded voltage indicators have also recently begun to show promising use *in vivo*^37^. Combining the temporal resolution of voltage transients with the spatial resolution of one-photon wide-field miniscope recording creates a powerful tool for imaging the brain during behavior, but such a technique is not without major caveats. Arguably, the biggest obstacle to incorporating voltage imaging in behavioral experiments is in the hardware capacity of the miniscope cameras and storage capacity of the recording computer. A miniscope CMOS capable of recording at several hundreds to a thousand frames per second would be needed to accurately capture high frequency spiking events, which would also result in much larger files sizes for storage and analysis. If anything, these challenges demonstrate a vast potential for growth and development for this research platform, with goals that are limited only by the capacity of existing technologies to study the brain.

### Long-term outlook and call for users

The ultimate and long-term goal of the M3 platform is to establish an extensive and freely accessible open research database mapping the functional neurodynamic qualities of the rodent and marmoset brains in multiple disease models and spanning a massive variety of standardized behavioral tasks. Such a database would grant neuroscientists around the world the freedom to browse through and download data from both wildtype and various disease models to analyze and compare population dynamics between brain regions and between species across a set of standardized cognitive tasks. Navigating the brain through this online atlas would greatly reduce the number of animals used in neuroscience research and would provide a valuable resource to computational researchers interested in conducting complex analyses between animal models tested across multiple cognitive modalities, who would need not bear the time and costs associated with purchasing and setting up their own *in vivo* recordings.

A collection of publicly accessible data of this scale would be impossible without the collaborative efforts of scientists within and beyond our institution, and we see great promise in extending this platform across multiple sites in the near future. To this end, we have started to assist several laboratories to setup their own miniscope/touchscreen recording systems and hope to expand these efforts going forward. Those interested in developing this platform at their home institutions are encouraged to contact the corresponding authors of this manuscript. In the same way that many connectome projects strive to map the network wiring of the brains of various living organisms, our hope is to provide an analogous atlas to single cell functionality in circuits throughout the brain to bring the field of neuroscience one big step closer to understanding the neural dynamics that underlie behavior and cognition in both healthy and diseased states.

